# TPM3.1 association with actin stress fibers is required for lens epithelial to mesenchymal transition

**DOI:** 10.1101/774794

**Authors:** Justin Parreno, Michael B. Amadeo, Elizabeth H. Kwon, Velia M. Fowler

**Affiliations:** Department of Molecular Medicine, The Scripps Research Institute, La Jolla, California; Department of Biological Sciences, University of Delaware, Newark, Delaware

## Abstract

**Purpose:** Epithelial to mesenchymal transition (EMT) is a cause of anterior and posterior subcapsular cataracts. Central to EMT is the formation of actin stress fibers. Targeting specific, stress fiber associated tropomyosin in epithelial cells may be a means to prevent stress fiber formation and repress lens EMT.

**Methods:** We identified Tpm isoforms in mouse immortalized lens epithelial cells and isolated whole lenses by semi-quantitative PCR followed Sanger sequencing. We focused on the role of one particular tropomyosin isoform, Tpm3.1, in EMT. To stimulate EMT, we cultured cells or native lenses in TGFβ2. To test the function of Tpm3.1, we exposed cells or whole lenses to a Tpm3.1-specific chemical inhibitor, TR100, as well as investigated lenses from Tpm3.1 knockout mice. We examined stress fiber formation by confocal microscopy and assessed EMT progression by αsma mRNA (qPCR) and protein (WES immunoassay) analysis.

**Results:** Lens epithelial cells express eight tropomyosin isoforms. Cell culture studies showed that TGFβ2 treatment results in an upregulation of Tpm3.1, which associates with actin in stress fibers. TR100 prevents stress fiber formation and reduces αsma in TGFβ2 treated cells. We confirmed the role of Tpm3.1 in lens epithelial cells in the native lens *ex vivo*. Culture of whole lenses in the presence of TGFβ2 results in stress fiber formation at the basal regions of epithelial cells. Knockout of Tpm3.1 or treatment of lenses with TR100 prevents basal stress fiber formation and reduces epithelial αsma levels.

**Conclusion:** Targeting specific stress fiber associated tropomyosin isoform, Tpm3.1, is a means to represses lens EMT.

## Introduction

The transition of lens epithelial cells into myofibroblasts through the process of epithelial to mesenchymal transition (EMT) is a cause of anterior subcapsular and posterior capsular cataracts. In EMT, myofibroblasts synthesize excessive fibrous matrix and contractile molecules ultimately leading to loss of lens architecture and light occlusion.

The reorganization of the actin into stress fibers is central to EMT. Stress fibers are thick bundles of polymerized actin that not only regulates migration, contraction, and matrix remodeling, but also influences the expression of myofibroblast molecules, such as αsma^1, 2^. Thus, targeting actin stress fibers may be a means to repress EMT. How to specifically target actin stress fibers during lens EMT remains unclear.

The tropomyosins (Tpms) are a family of proteins that bind and stabilize actin filaments (F-actin). The Tpms are highly conserved with over forty non-redundant mammalian isoforms generated from alternative splicing of four Tpm genes: Tpm1, 2, 3, and 4. Each isoform has a unique interaction with actin filaments as well as a specific ability to coordinate the interaction of other binding proteins to actin. Thus, the specific expression pattern of Tpms within a cell defines actin organization^3^.

Several Tpms associate with actin stress fibers ^4-9^. In osteosarcoma (U2OS) cells, there are six known stress fiber-associated Tpm isoforms: Tpm1.6, Tpm1.7, Tpm2.1, Tpm3.1, Tpm3.2 and Tpm4.2 ^8, 9^. Each isoform plays a distinct role in regulating actin stress fibers by coordinating interactions with specific actin binding proteins such as non-muscle myosins, actin severing proteins, and/or actin nucleating proteins. Elimination of any of stress fiber associated Tpm can compromise stress fiber formation^8^. While previous studies in the lens have implied a role of Tpms from the Tpm1 and 2 gene in cataract formation^10, 11^, an understanding on the specific isoforms expressed and their participation in stress fibers formation by lens epithelial cells has not been shown. Here, we systematically identified all the Tpm isoforms in mouse lens epithelial cells associated with stress fiber formation and showed that Tpm3.1 is the most highly upregulated isoform upon stimulation with TGFβ. Inhibition of Tpm3.1 with a small molecule chemical inhibitor, or by genetic deletion revealed that Tpm3.1 plays a critical role in stress fiber formation during EMT in the mouse lens. Our strategy to identify and inhibit stress fiber-associated Tpms may be broadly applicable in targeting stress fibers to repress EMT in cataracts.

## Materials and Methods

### Cell culture

Immortalized mouse lens epithelial cells (imLECs) ^12^ were a gift from Dr. Xiaohua Gong (University of California Berkeley). ImLECs were routinely passaged in growth media, which consisted of Dulbecco’s Modified Eagle’s medium (DMEM) supplemented with 10% fetal bovine serum (FBS) and 1% penicillin-streptomycin, at 37°C and 5% CO_2_. Cells were cultured until 70-90% confluency at which point cells were dissociated from culture vessels using 0.05% Trypsin-EDTA (Thermo Fisher Scientific; Waltman, MA) and reseeded onto culture vessels for subsequent passaging. Cells were maintained up to 15 passages.

Cells were seeded at an approximate density of 2.5 × 10^4^ cells/cm^2^ onto either tissue culture polystyrene or glass-bottomed dishes in complete media for experiments. After 24 hours, the cells were placed in serum-reduced media (DMEM supplemented with 0.5% FBS). After an overnight incubation, the media was replaced with fresh serum-reduced media supplemented with 5ng/mL TGFβ2 in the presence or absence of TR100.

### Cell viability assays

Following 24 hours of TR100 treatment, cell viability was measured. Adherent imLECs were detached from culture dishes using 0.05% Trypsin-EDTA and then centrifuged. After pelleting, the cells were resuspended in fresh media, and an aliquot was mixed in a 1:1 ratio with Trypan blue. Cell counts were conducted on a TC20 automated cell counter (Bio-Rad). While cells treated with <5μM of TR100 remained adherent, treatment of imLECs with higher concentrations of TR100 lead to cell detachment. To count floating cells, the media containing the cells was harvested and centrifuged. Pelleted cells were resuspended in fresh media and an aliquot was mixed in a 1:1 ratio with Trypan blue. Approximately 94.1% and 100% were floating following treatment with 10μM or 20μM TR100, respectively. The large majority of floating cells (>87.4%) were not viable (Trypan blue positive). To calculate total percent viability, the number of live cells in floating and adherent fractions were added and then divided by total sum of live and dead cells in both fractions.

Cell viability was also assessed visually by staining cells cultured on glass dishes with 1μM of Calcein-AM (BioVision, Milipitas, CA) and TO-PRO3 (Thermo Fisher Scientific) to determine live and dead cells, respectively. Cells were stained for 20 minutes, followed by three washes in PBS, and then confocal fluorescence microscopy.

### Mice and lens dissections

Wild-type and *Tpm3/Δexon9d*^*-/-*^ mice between the ages of 8 and 10 weeks in the C57BL/6J background were used for experiments. The *Tpm3/Δexon9d*^*-/-*^ mouse strain which has been previously described^13^, was a generous gift from Dr. Peter Gunning (University of New South Wales). Mice were backcrossed to wild-type C57BL/6J to generate congenic strains. All animal procedures were conducted in adherence to the ARVO Statement for the Use of Animals in Ophthalmic and Vision Research and performed in accordance with approved animal protocols from The Scripps Research Institute and the University of Delaware.

Following euthanasia, eyes were enucleated and lenses were dissected as previously described^14^. Lenses were immediately placed in medium 199 supplemented with 1% penicillin-streptomycin. To stimulate EMT, lenses were placed in medium 199 (Life Technologies) supplemented with 5ng/mL TGFβ2 (Thermo Fisher Scientific).

### RT-PCR gene expression analysis and sequencing

To isolate RNA from imLECs, cells within a six-well plate were homogenized in 1mL of TRIzol reagent (Thermo Fisher Scientific) and stored at −80°C until further processing. To isolate RNA from lens epithelium, lens epithelium was separated from fiber cells. To achieve this, lens capsules were punctured at the lens equator using fine forceps. With a second set of fine tweezers, the lens capsules were peeled from the fiber cell mass. At least six pooled lens capsules, which contained mostly epithelial cells, were immersed in 300μL of TRIzol. In a separate tube, lens fiber masses from two pooled lenses were immersed in 300μL of TRIzol.

RNA from tissue or cells were then processed according to manufacturer’s instructions. Reverse transcription was performed on equal amounts of total RNA for each sample using Superscript III First-Strand Synthesis System for RT-PCR Kit (Thermo Fisher Scientific).

Semi-quantitative PCR was performed using Hot-Start Taq Blue Mastermix (Thomas Scientific, Swedesboro, NJ) on equivalent amounts of cDNA in a 20μL reaction and using previously validated primers^15^. Equivalent volumes of PCR product were loaded for electrophoresis on a 1% agarose gel. Under ultraviolet illumination, PCR bands were cut from gels. Products were purified using a gel extraction kit (QIAquick; Qiagen, Germantown, Maryland) and submitted for Sanger sequencing (Genewiz; La Jolla, CA).

Relative real-time PCR was performed using PowerSYBR Green PCR Master Mix (Thermo Fisher Scientific) in a 25uL reaction according to manufacturer’s directions, using a CFX-96 real-time system (Bio-Rad). qPCR primers were developed using NCBI primer-blast. Primers for αsma (forward, 5’-TCACCATTGGAAACGAACGC-3’; reverse, 5’-CCCCTGACAGGACGTTGTTA-3’) and Gapdh (forward, 5’-TTCGAGAGTCAGCCGCATTT-3’; reverse, 5’-ATCCGTTGACTCCGACCTTC-3’) were based on NCBI reference sequences NM_007392.3 and NM_001289726.1, respectively.

### Total protein extraction and WES capillary-based immunoassay

Cells were washed once in PBS and then harvested from culture by scraping into fresh PBS. Cells were pelleted by centrifugation at 800 x g for 5 minutes and protein was extracted by incubating pellets in 1x RIPA buffer (20mM Tris-HCL, pH 7.4, 150mM NaCl, 1mM Na_2_EDTA, 1mM EGTA, 1% NP-40, 1% sodium deoxycholate, 2.5mM sodium pyrophosphate, 1mM β-glycerophosphate, 1mM Na_3_VO_4_, and 1mg/mL leupeptin) with Protease Inhibitor Cocktail (1:100; P8430, Sigma-Aldrich).

To extract protein from lens epithelium, lens epithelia were first mechanically separated from fibers as above. Each lens epithelium was placed in 10μL of 1x RIPA buffer with Protease Inhibitor Cocktail. Extracts were incubated on ice for 30 minutes and then briefly sonicated. Protein was quantified using bicinchoninic acid (BCA) protein assay (Thermo Fisher Scientific). Equal amounts of protein were prepared for capillary-based (Wes) immunoassay using a 12-230 Separation module kit according to manufacturer’s instructions (Protein Simple, San Jose, CA). The concentration of sample protein and antibodies used in experiments were in the dynamic range of protein level detection and was determined through empirical optimization. Total protein of samples loaded into capillaries was determined using a Total Protein Separation module kit (Protein Simple). Protein expression of specific molecules were normalized to total protein.

### Triton X-100 fractionation of imLECs

Triton X-100 soluble and insoluble proteins were separated as previously described with slight modifications^16, 17^. Briefly, imLECs cultured in 6-well dishes were washed once in PBS and then incubated in 72μL of Triton-extraction buffer (0.1% Triton X-100, 1mM EGTA, 10mM PIPES, 100mM NaCl, 3mM MgCL_2_, 300mM sucrose) for 2 minutes at room temperature. The Triton-extracted soluble portion was then pipetted into a 1.5 ml Eppendorf tube. Approximately 8μL of 10x RIPA was added then to the solution. The remaining Triton-insoluble mLEC fractions were isolated by directly scraping into 80μL of 1x RIPA buffer and pipetted into a separate 1.5 ml Eppendorf tube. Equal volumes were prepared for Wes immunoassay.

### Immunostaining of imLECs

Cells cultured on glass-bottom dishes (World Precision Instruments, Sarasota, FL) were fixed in 4% paraformaldehyde (Electron Microscopy Sciences, Hatfield, PA) at room temperature for 10 minutes. Samples were then washed in PBS and kept at 4°C until further processing. For immunostaining cells were permeabilized for 30 minutes in permeabilization/blocking solution (PBS containing 0.3% Triton, 0.3% bovine serum albumin, 3% goat serum) and then incubated with mouse anti-tropomyosin-5 antibody (1:200; clone 2G10.2, Millipore, Burlington, MA) which recognizes Tpm3.1 and Tpm3.2. Following an overnight incubation at 4°C, cells were washed three times in PBS, 5 minutes per wash, and then incubated at room temperature for 1 hour in secondary antibody solution which contained Alexa 488-conjugated anti-mouse antibody (1:200; Thermo Fisher Scientific), rhodamine phalloidin (1:50; Thermo Fisher Scientific), and Hoechst 33342 (1:500; Thermo Fisher Scientific). Cells were washed and then mounted using ProLong gold anti-fade reagent (Thermo Fisher Scientific).

### Immunostaining of native lens epithelial cells

For immunostaining of native lens epithelial cells, whole lenses were fixed by immersing lenses in 4% paraformaldehyde in PBS at room temperature for 30 minutes. Fixed lenses were then washed three times in PBS (5 minutes per wash). To allow for penetration of antibody, each lens was immersed in 0.2mL (0.25%) collagenase A (Roche Diagnostics, Mannheim, Germany) solution (in PBS) in individual wells of a 48 well plate and incubated at 37°C. After 20 minutes of collagenase digestion, lenses were washed (5 minutes per wash) three times in PBS and placed in permeabilization/blocking solution at room temperature for 2 hours. Next, lenses were labeled with primary mouse anti-tropomyosin-5 antibody (1:100; clone 2G10.2) in permeabilization/blocking solution and incubated at 4°C overnight. Lenses were then washed three times in PBS and placed in permeabilization/blocking solution containing Alexa 488-conjugated anti-mouse secondary antibody (1:200), CF640 dye-conjugated to wheat germ agglutinin (WGA) (Biotium, Fremont, CA) (1:100), rhodamine-phalloidin (1:20), and Hoechst 33342 (1:500) at room temperature for 2 hours. Lenses were washed three times in PBS for 5 minutes prior to imaging by confocal fluorescence microscopy.

### Confocal fluorescence microscopy

Imaging was performed using a Zeiss 880 laser-scanning confocal fluorescence microscope (Zeiss, Germany) using a 20× 0.8 NA objective, a 63× 1.4 NA oil objective, or a 100× 1.4 NA oil objective. Raw images were processed using Zen (Zeiss) software. Image line scan analysis was performed using FIJI software.

### Statistical analysis

Each experiment was replicated on at least three separate occasions. Statistical analysis was performed using GraphPad Prism7 (San Diego, CA). Unpaired *t* tests were used to detect differences between two groups of data. To determine differences between multiple groups of data, an analysis of variance followed by post-hoc testing was performed. A Dunnett’s post-hoc test was used to compare differences between several experimental groups against a control. A planned comparisons test with a Sidak’s adjustment post-hoc was used to measure differences between multiple groups of data.

## Results

### Tpm expression in immortalized lens epithelial cells

To determine Tpm isoform expression in imLECs, we performed semi-quantitative RT-PCR using previously validated primer sets^15^. The cDNA products were extracted from agarose gels and were sequenced. We determined that imLECs express eight Tpm isoforms (Figure 1A): Tpm 1.5, 1.7, 1.9, 1.13, 2.1, 3.1, 3.5, and 4.2.

**Figure 1.**
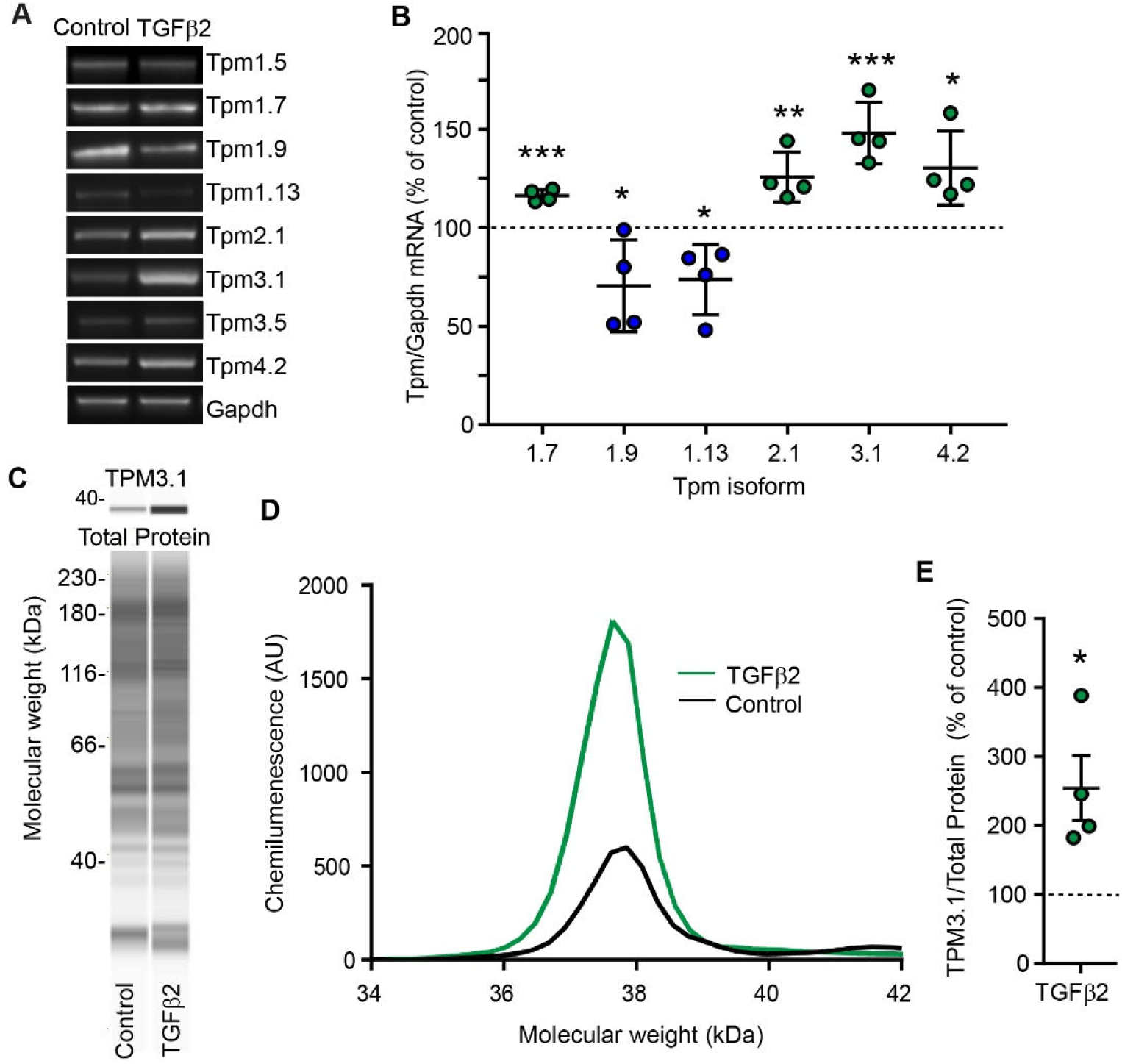
Lens epithelial cells express eight Tpm isoforms including Tpm 3.1, a stress fiber associated Tpm, that is upregulated by TGFβ2 treatment. (A) Semi-quantitative RT-PCR for Tpms in imLECs treated with TGFβ2 for 1 day. The bands in gels were cut to extract cDNA in order to confirm transcript identity by gene sequencing. (B) TGFβ2 modulates specific Tpm mRNA levels by downregulation Tpm1.9 and 1.13. TGFβ2 also upregulates known stress fiber associated Tpms: Tpm1.7, 2.1, 3.1 and 4.2. TGFβ2 most highly upregulates Tpm3.1. WES capillary-based immunoassay demonstrates increases in Tpm3.1 protein levels. Panel (C) shows a generated pseudo-Western blot of Tpm3.1 and total protein. (D) is an electropherogram of Tpm3.1 and (E) represents a dot-plot of cumulative data points for Tpm3.1 protein levels following TGFβ2 treatment.

Next, we induced EMT by exposing imLECs to TGFβ2. We chose to treat cells with TGFβ2 as it has been shown to be present in the aqueous humor ^18^, it is increased in the aqueous humor following lens injury and inflammation ^19, 20^, and it has been shown to be more potent than TGFβ1 in enhancing EMT ^21^. Exposure of imLECs to 5ng/mL TGFβ2 downregulated Tpm 1.9 and Tpm1.13 mRNA levels, upregulated Tpm1.7, 2.1, 3.1, and 4.2 mRNA levels, but did not affect Tpm1.5 and Tpm3.5 mRNA levels (Figure 1A and 1B).

Tpm1.7, 2.1, 3.1, and 4.2 have been shown to associate with actin in stress fibers in U2OS cells^8, 9^. Since elimination of any of these Tpms can compromise stress fiber formation^8^, we chose to further examine the role of Tpm3.1, the most highly upregulated Tpm isoform in imLECs by TGFβ2 treatment. Using WES immunoassay, we confirmed that TGFβ2 upregulates Tpm3.1 protein level (Figure 1C-E).

### A pharmacological Tpm inhibitor, TR100, reduces the assembly of Tpm3.1 into F-actin stress fibers

To elucidate the function of Tpm3.1, we exposed imLECs to TR100. TR100 is a Tpm inhibitor that associates with a binding pocket located in the 9D exon of Tpm3.1 and prevents end-to-end assembly of Tpm, reducing Tpm binding along actin filaments in cells^22^. To determine the optimal concentration of TR100, we assessed cell viability at different concentrations of TR100. Treatment of imLECs with 5μM or 10μM TR100 causes cell death, however, treatment with 1μM or 2μM of TR100 does not significantly impact imLEC viability (Figure 2A, B). Based on these findings, we chose to use 1μM of TR100 for subsequent experiments.

**Figure 2.**
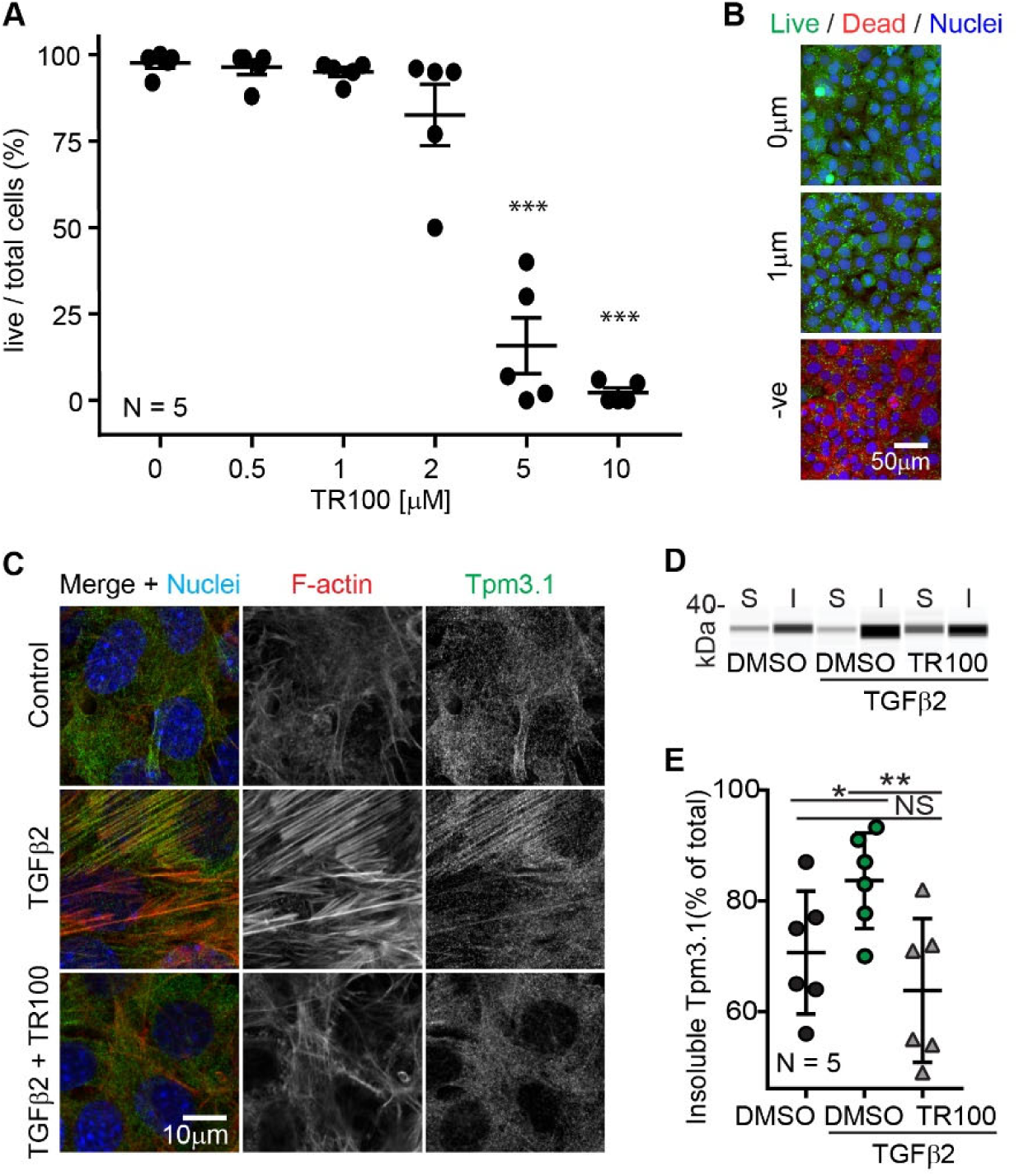
1μM TR100 treatment reduces the induction of actin stress fibers by TGFβ2. (A) Dot plot showing the percentage of live cells following treatment with different concentrations of TR100. (B) imLECs are viable following treatment with 1μM of TR100 for 1 day as seen by positive staining for calcein AM staining (live cells; green) and negative staining for TO-PRO-3 (dead cells; red). Cells fixed in 4% paraformaldehyde are used as negative, dead controls. (C) Immunostaining of imLECs for F-actin (red) and Tpm3.1 (green). TGFβ2 treatment leads to association of Tpm3.1 with F-actin and actin stress fibers formation. TR100 treatment reduces Tpm3.1 association with F-actin and represses stress fiber formation. (D) WES capillary assay pseudo-Western blot and corresponding (E) dot-plot showing an increase in Tpm3.1 in the cytoskeletal (Triton insoluble) fractions of imLECs.

To determine if Tpm3.1 inhibition prevents stress fiber assembly, we exposed imLECs to TGFβ2 supplemented with TR100. As expected, treatment of imLECs with TGFβ2 results in prominent F-actin assembly into stress fibers that are associated with Tpm3.1 (Figure 2C). By contrast, TR100 treatment reduces the formation of stress fibers in TGFβ2-treated imLECs along with substantially reduced association of Tpm3.1 along F-actin stress fibers. As a confirmation of our immunocytochemistry findings, we used WES immunoassay to examine the proportion of Tpm3.1 protein co-sedimenting with the F-actin cytoskeleton after Triton-X100 extraction. This reveals that Tpm3.1 is present in both Triton-soluble and insoluble fractions of untreated cells (Figure 2D and 2E) and that treatment with TGFβ2 results in a greater proportion of Tpm3.1 in the Triton insoluble fraction, consistent with enhanced Tpm3.1 assembly with F-actin into stress fibers. Similar to the immunofluorescence staining results (Figure 2C), treatment with TR100 reduces the proportion of Triton-insoluble Tpm3.1 in TGFβ2-treated cells (Figure 2D, E). We conclude that selective inhibition of Tpm3.1 function impairs TGFβ2-induced stress fiber formation in imLECs.

### TR100 represses the transition of imLECs into myofibroblasts

To determine if inhibition of Tpm3.1 could prevent EMT progression in imLECs, we examined the expression of αsma, the hallmark of myofibroblasts^23, 24^. Relative real-time RT-PCR demonstrates that TGFβ2 treatment enhances αsma mRNA levels 2.3-fold over untreated cells (Figure 3A). These increases in αsma mRNA levels can be repressed completely by inclusion of with TR100 during the TGFβ2 treatment. Furthermore, WES immunoassay (Figure 3B-D). indicates that TGFβ2 treatment enhances αsma protein levels 5.7-fold, an increase which is nearly completely repressed by TR100 treatment. Thus, TR100 represses both αsma mRNA and protein levels in TGFβ-treated imLECS and reduces EMT based on this marker.

**Figure 3.**
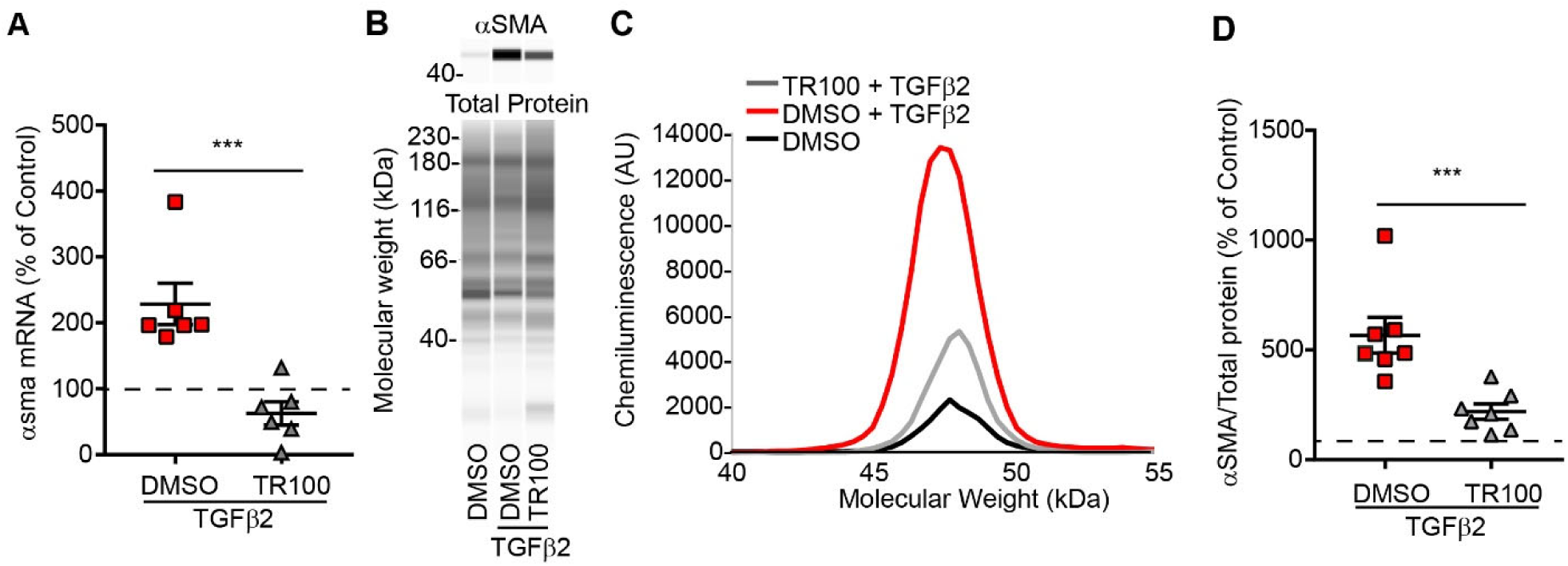
TR100 reduces αSMA expression in imLECs. (A) Relative qRT-PCR demonstrates that TR100 treatment represses the induction of αsma mRNA levels by TGFβ2 after 1 day treatment. Wes capillary assay (B) pseudo-Western blot, (C) electropherogram, and (D) corresponding dot-plot showing that TR100 treatment represses the induction of αSMA protein levels after 2 days.

### Lens epithelial cells express eight Tpm isoforms including Tpm 3.1

To characterize the role of Tpm3.1 in a physiological model, we turned to whole native lenses. First, we determined Tpm isoform expression in lens epithelial cells and compared to Tpm isoform expression in fiber masses, after separating lens epithelia from fiber masses prior to RNA extraction. Since there is approximately 6.3-fold less RNA in lens epithelium as compared to fiber masses (data not shown), we reverse transcribed an equivalent amount of RNA from lens epithelium and fibers and then performed semi-quantitative PCR followed by Sanger sequencing of cDNA products. Native lens epithelial cells express eight Tpm isoforms (Figure 4A). Five of the isoforms (Tpm1.5, 1.7, 1.9, 2.3, 3.1) are detected solely in the lens epithelium but not in fiber cells, whereas three of the isoforms (Tpm1.13, 3.5, and 4.2) are expressed in both the lens epithelium and fiber cells. Tpm isoform expression of native lens epithelial cells differs from the imLECs by only one isoform; native lens epithelial cells express Tpm2.3 (Figure 4A) whereas imLEC s express Tpm2.1 (Figure 1A). Of note, Tpm3.1 is expressed in native lens epithelial cells but not in lens fiber cells. The faint cDNA band that appears in gels of the Tpm3.1 RT-PCR reaction for fiber cells is not of Tpm3.1 based on Sanger sequencing.

**Figure 4.**
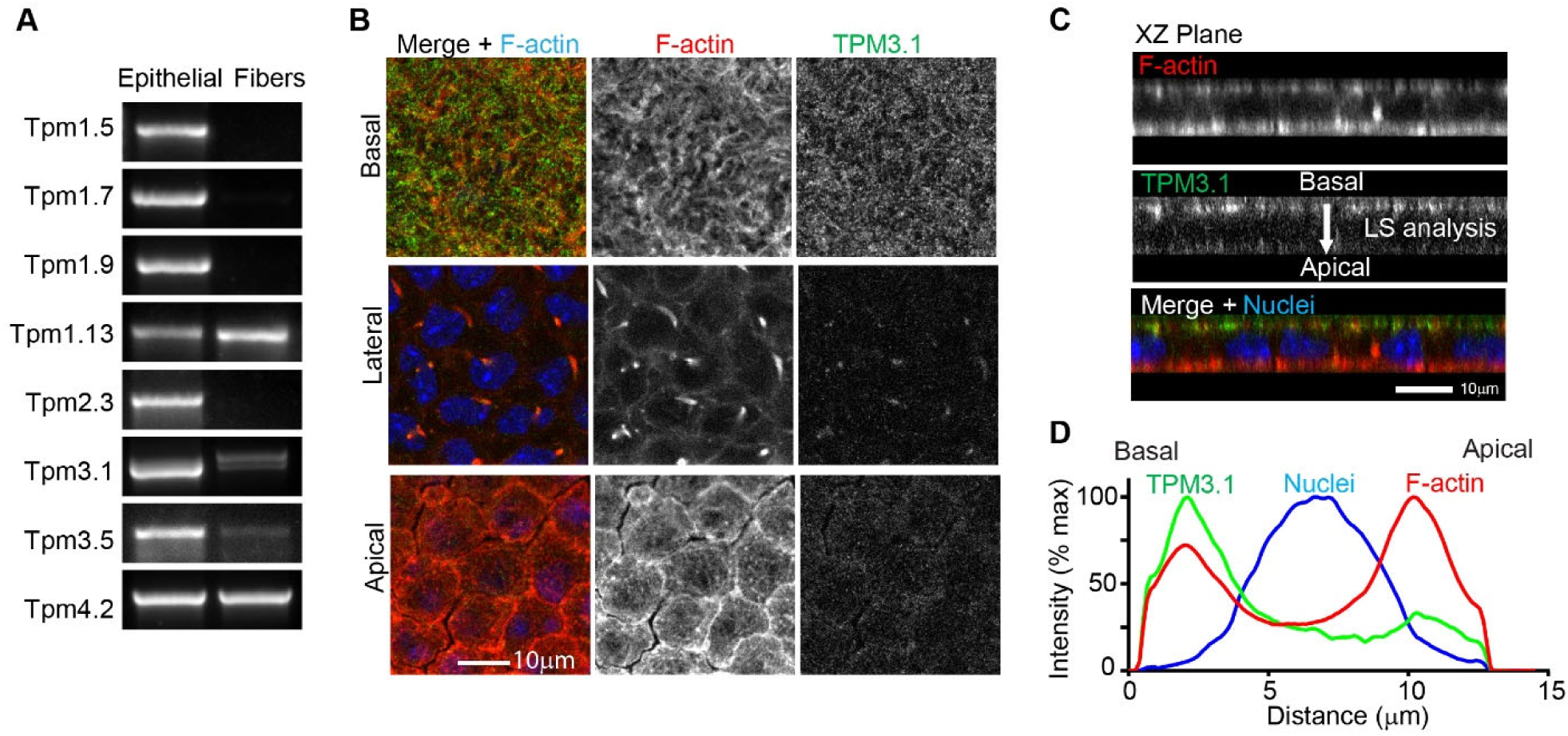
Tpm3.1 is expressed in native lens epithelial cells and associates with basal F-actin. (A) Semi-quantitative PCR on equivalent amounts of cDNA from epithelial and fiber cell RNA reveals the expression of eight Tpms in native lens epithelial cells, including the expression of Tpm3.1. (B) Single (*x,y*-plane view) optical sections of basal, lateral, and apical regions of the lens showing F-actin organization (phalloidin; red), Tpm3.1 localization (green) and nuclei (Hoecsht; blue). (C) Sagittal (*x,z*-plane view) optical sections from 3D reconstructions of confocal z-stacks showing Tpm3.1 (green) associates with F-actin (phalloidin; red) at the basal regions of cells. This was confirmed by (D) line scan analysis of images.

To determine the presence and localization of Tpm3.1 protein within native lens epithelial cells, we developed a methodology for fluorescent immunostaining of proteins within lens epithelial cells. Using this methodology, we can image Tpm3.1 in epithelial cells of native lenses (Figure 4B, C). The intensity of Tpm3.1 staining is greatest at the basal regions of lens epithelial cells with very little staining at the middle and apical regions. At the basal regions, there were no apparent stress fibers, and Tpm3.1 is localized in a punctate pattern partially co-localized with F-actin (Figure 4B). XZ plane reconstruction of confocal images showed that while F-actin staining is most intense in the apical region of epithelial cells, Tpm3.1 staining is most intense at the basal region of cells (Figure 4C). We confirmed through a basal-to-apical line scan analysis of fluorescent intensity (Figure 4D).

### Tpm3.1 knockout or TR100 treatment represses stress fiber formation in lens epithelial cells of native lenses

Next, we developed an *ex vivo* culture model for EMT using whole lenses^21, 25^. Whole organ culture of lenses is a useful tool to study lens EMT as the overall native three-dimensional architecture of the lens is preserved ^21, 26^. In this model, to stimulate EMT, we cultured mouse lenses in medium 199 in the absence and presence of TGFβ2 and compared to non-cultured (control) lenses. After two days of culture, we determined the cellular region where F-actin stress fibers form during lens epithelial cell EMT using whole mount confocal imaging of lenses stained with phalloidin. The culture of lenses led to stress fiber formation at the basal regions of lens epithelial cells which are absent in control lenses (Figure S1; Figure 4B). Since culture of lenses in medium 199 supplemented with TGFβ2 results in greater stress fiber formation than culture of lenses in medium 199 alone, to stimulate EMT in subsequent studies, we cultured lenses in medium 199 supplemented with TGFβ2.

To study the role of Tpm3.1, we examined the actin stress fibers at the basal regions of lens epithelial cells in Tpm3.1 knockout (*Tpm3/Δexon9d*^*-/-*^) mice. Freshly fixed lenses *Tpm3/Δexon9d*^*-/-*^ had similar actin organization to wildtype, *Tpm3/Δexon9*^*+/+*^ mice (Figure 5; Figure S2). Notably, there is an absence of actin stress fibers at the basal regions of the epithelial cells (Figure 5). Exposure of wildtype, *Tpm3/Δexon9*^*+/+*^ to TGFβ2 led to formation of actin stress fibers as shown above (Figure S1). However, stress fiber formation at the basal region of epithelial cells in TGFβ2-treated *Tpm3/Δexon9d*^*-/-*^ lenses is repressed as compared to *Tpm3/Δexon9*^*+/+*^ lenses. F-actin organization at the middle and apical region of *Tpm3/Δexon9*^*+/+*^ and *Tpm3/Δexon9d*^*-/-*^ lens epithelial cells had similar organization following TGFβ2 treatment (Figure S2). Thus, knockout of Tpm3.1 specifically represses the effect of TGFβ2 on F-actin stress fiber formation at the basal region of epithelial cells.

**Figure 5.**
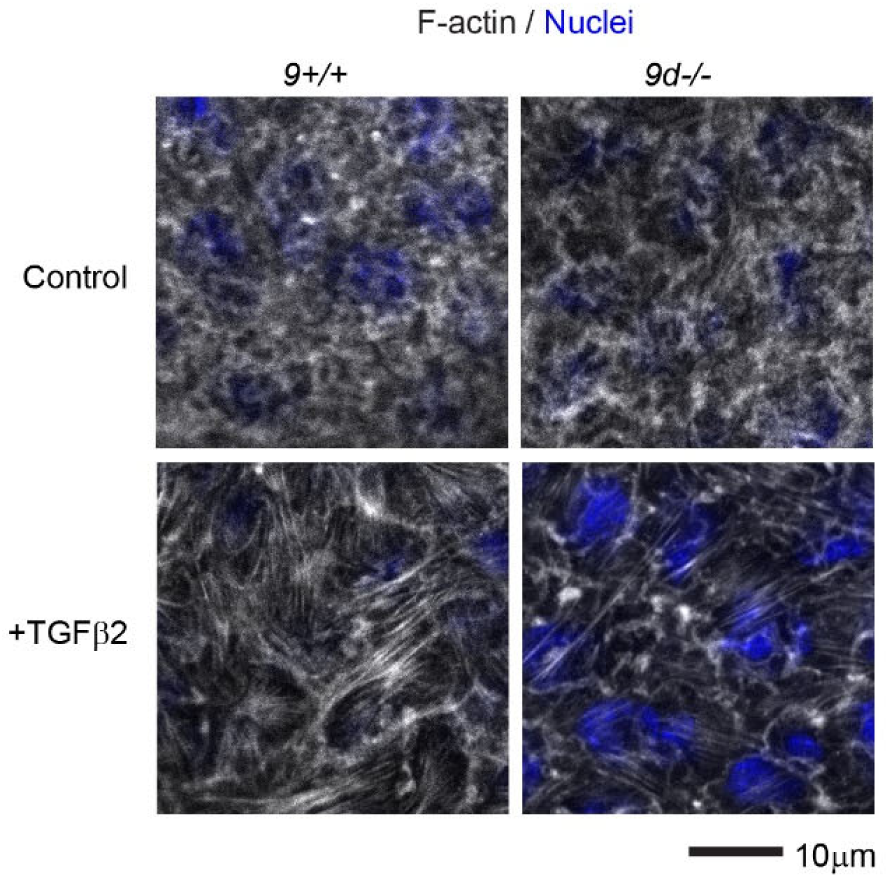
Stress fiber formation is repressed in *Tpm3/Δexon9d*^*-/-*^ lenses. Whole mount images showing single (*x,y*-plane view) optical sections of basal regions of the lens epithelium in non-cultured and lenses cultured with TGFβ2 for 2 days. In non-cultured lenses, F-actin staining (phalloidin; grayscale) is similar in *Tpm3/Δexon9d*^*-/-*^ (9d-/-) lenses as compared to *Tpm3/Δexon9*^*+/+*^ (9d+/+) lenses. However, stress fiber formation at the basal epithelial regions is repressed in *Tpm3/Δexon9d*^*-/-*^ lenses as compared with *Tpm3/Δexon9d*^*-/-*^ lenses following TGFβ2 treatment.

To determine if the Tpm3.1 inhibitor TR100 also prevents stress fiber assembly in native lens epithelial cells, we exposed whole lenses to TGFβ2 in the presence of 1μM TR100. Similar to our results with imLECs (Figure 2C), TR100 reduces F-actin stress fiber formation in the basal region of native lens epithelial cells (Figure 6A) and reduces the association of Tpm3.1 with F-actin in the stress fibers (Figure 6B). Reduction of TGFβ2-induced stress fibers both in absence of Tpm3.1 in *Tpm3/Δexon9*^*-/-*^ lenses, or in TR100-treated wild-types lenses provide evidence that normal levels and function of Tpm3.1 is essential for stress fiber formation at the basal region of lens epithelial cells.

**Figure 6.**
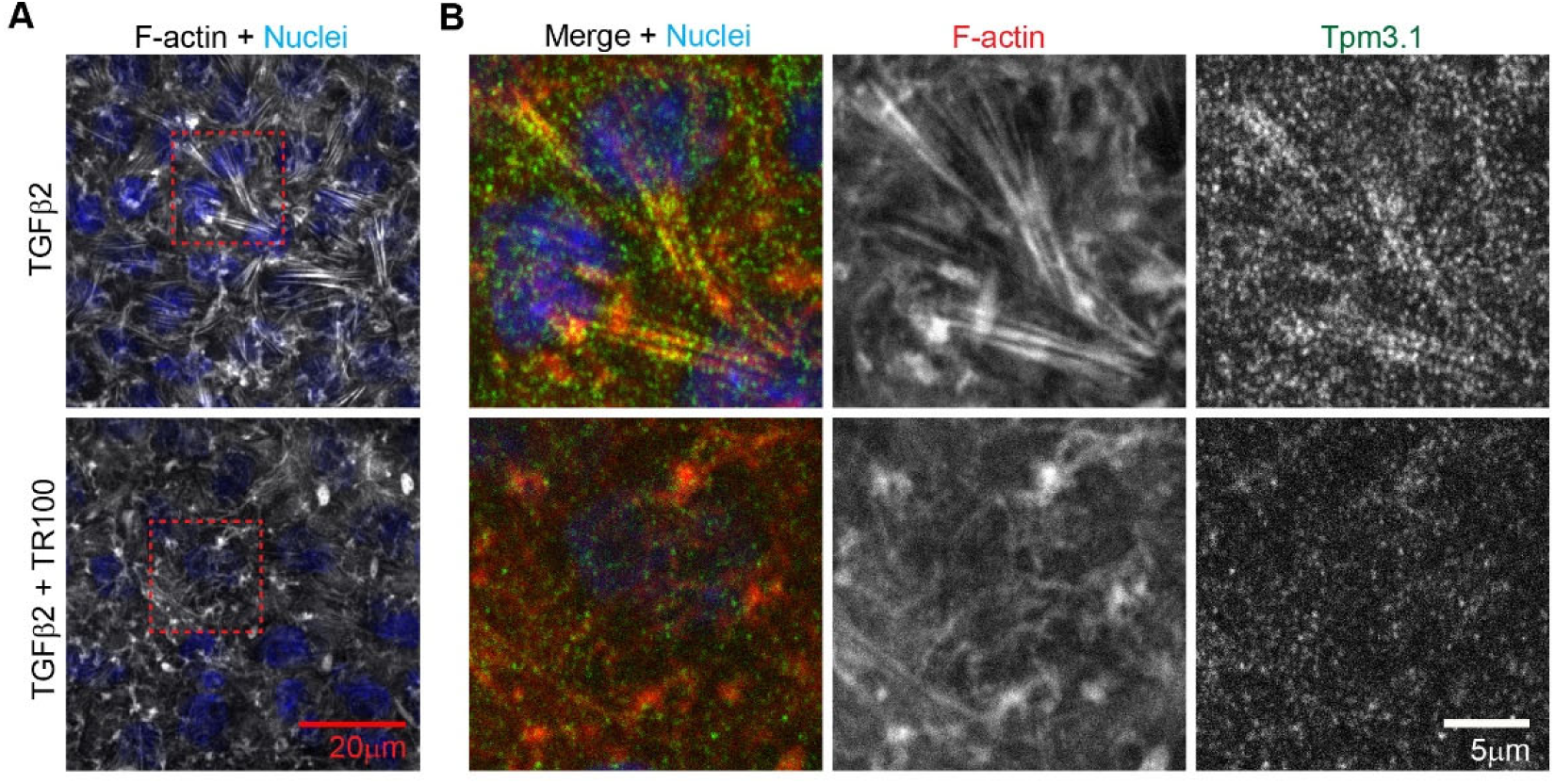
Stress fiber formation at the basal regions of native lens epithelial cells is repressed by TR100. Whole mount images of (A) single (*x,y*-plane view) optical section of the basal region of lens epithelial cells revealing that stress fiber formation is reduced in TR100 treated lenses for 2 days. (B) Tpm3.1 (green) immunostaining demonstrates association with F-actin (red) in stress fibers. TR100 treatment reduces the association of Tpm3.1 with F-actin.

### The induction of αsma by TGFβ2 is repressed in Tpm3/Δexon9^-/-^ lenses or by treatment of wild-type lenses with TR100

To determine if knockout of Tpm3.1 or inhibition with TR100 reduces EMT progression into myofibroblasts, we examined αsma protein levels. For this analysis we extracted protein from capsular-epithelial peels. By performing WES immunoassay, which is a sensitive means to detect protein in a low volume of extract, we can determine αsma protein expression from individual peels (Figure 7). We determined that TGFβ2 treatment led to 2.8-fold increase in αsma protein levels as compared to control. Knockout of Tpm3.1 represses (*Tpm3/Δexon9*^*-/-*^ lenses) represses this increase in αsma levels. In addition, exposure to TR100 also reduces αsma levels in TGFβ2 treated wildtype lenses. Therefore, knockout or inhibition of Tpm3.1 represses EMT progression in native lens epithelial cells.

**Figure 7.**
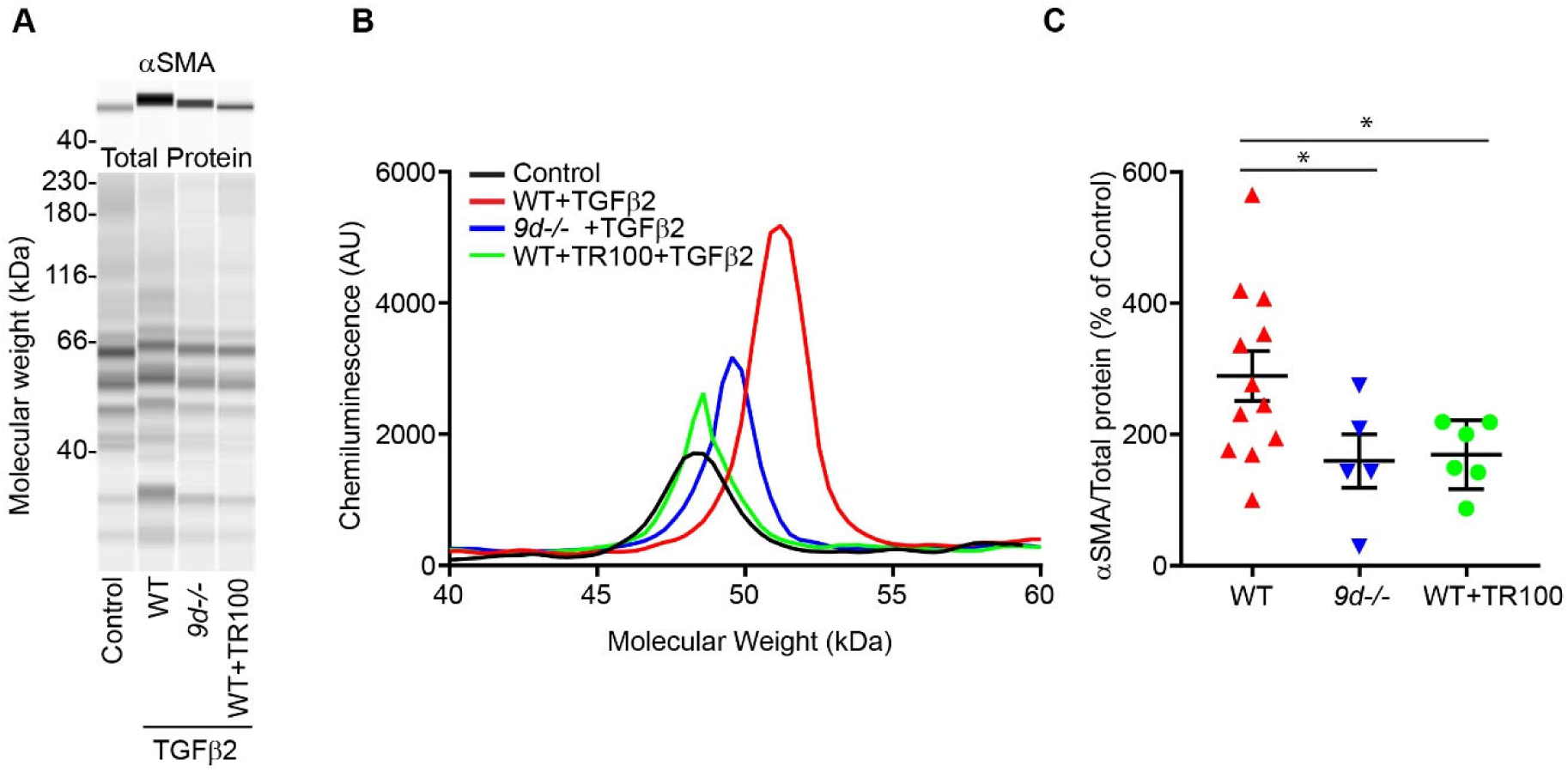
Knockout or inhibition of Tpm3.1 reduces αSMA expression in native lens epithelial cells. (A) pseudo-Western blot, (B) electropherogram, and (C) corresponding dot-plot showing αSMA protein levels in wild-type (WT; Tpm3/Δexon9d^+/+^) or knockout (9D^-/-^; Tpm3/Δexon9d^-/-^) mouse lens epithelial peels. Lens epithelial peels were from lenses either control (harvested fresh), cultured for four days with TGFβ2, or cultured for four days with TGFβ2 in the presence of TR100.

## Discussion

This study provides new insights into targeting stress fiber formation via inhibition of specific tropomyosin isoforms during lens EMT. We found four Tpm isoforms are upregulated in imLECS following TGFβ2 treatment, including Tpm3.1 which is the most highly upregulated. Tpm3.1 is expressed in imLECs and native lens epithelial cells but not fiber cells. During TGFβ2 induced EMT in imLECs or whole lenses, Tpm3.1 associates with F-actin in stress fibers. Knockout of Tpm3.1 or inhibition with the Tpm3.1-binding molecule TR100 prevents stress fiber formation and EMT progression in lens epithelial cells.

Tpm3.1 is the predominantly upregulated Tpm isoform in imLECS following TGFβ2 treatment and is associated with F-actin in stress fibers. The association of Tpm3.1 with actin stress fibers in lens epithelial cells is in agreement with previous studies on osteosarcoma (U2OS)^8, 27^, neuroblastoma (SK-N-BE(2))^28^ and neuronal (B35) cells^5^. Tpm3.1 enhances non-muscle myosin IIA phosphorylation and recruitment of non-muscle myosin IIA to stress fibers^5, 8^. Furthermore, Tpm3.1 can stimulate the activation of myosin II ATPase ^9^, and the duty ratio of non-muscle myosin IIB on F-actin ^28^ presumably leading to enhanced contractility. Tpm3.1 can also protect against F-actin severing by cofilin/actin-depolymerization factor^5, 29^ but also promote actin severing via gelsolin (KD = 0.3 ± 0.2 μM)^30^. Actin severing via gelsolin results in the creation of free filament ends and an overall net increase in actin polymerization by actin monomer addition. Thus, Tpm3.1 can promote stress fiber formation and actomyosin contractility in lens epithelial cells through several possible mechanisms.

In addition to Tpm3.1, we found Tpm1.7, 2.1, and 4.2 to be upregulated in imLECs by TGFβ2 treatment. These Tpms are also reported to associate with actin stress fibers and play diverse roles in promoting stress fibers^8, 9, 27^. Tpm1.7 protects against actin severing by cofilin^9^ and can promote the incorporation of αsma into stress fibers^31^. Tpm4.2 activates myosin II ATPase and is present mostly in highly bundled F-actin^9^. Furthermore, Tpm2.1 can promote actin filament elongation^32^. Stress fiber formation in Tpm3.1 knockout lenses or in lenses treated with TR100 is impaired, but not completely abolished. Thus, while impairment of Tpm3.1 may compromise stress fiber formation, the other Tpms may also promote stress fiber formation to some extent. Further research into the roles of other stress fiber-associated Tpms in lens EMT is required.

Furthermore, we found that Tpm3.1 is expendable for native lens epithelial actin organization, as freshly isolated knockout lenses had similar F-actin organization as wildtype lenses (Figure S2). F-actin organization at cell-cell junctions (middle and apical regions) and in polygonal arrays (apical region) were similar in *Tpm3/Δexon9*^*+/+*^ and *Tpm3/Δexon9d*^*-/-*^ lenses. This suggests that other Tpms may be more important for establishing F-actin organization in native lens epithelial cells. For example, lens epithelial cells contain Tpm 1.9, which has been shown to be associated with F-actin at adherens junctions in other (LLC-PK1 kidney) epithelial cells^33^. Tpm1.9 may play a similar role in lens epithelial cells based on several observations. In contrast to the stress fiber-associated Tpms, TGFβ2 downregulates Tpm1.9 expression. This corresponds with TGFβ2-induced decreases in F-actin levels at the adherens junctions in lateral and apical regions of the epithelial cells, which was not prevented in the absence of Tpm 3.1. Thus Tpm1.9 may be essential for maintaining F-actin at the adherens junctions. This also suggests that complete prevention of EMT may require increases in levels or activities of other Tpms such as Tpm1.9. Of note, Tpm1.13 is also downregulated by TGFβ2 treatment. Further studies are required to determine the role of both Tpm1.9 and 1.13 in epithelial cells. Targeting Tpm1.9 and 1.13 to promote their F-actin binding activities, may be required to fully repress lens EMT.

In conclusion, lens epithelial express several Tpm isoforms. Inhibition of stress fiber-associated Tpms, such as Tpm3.1, may be a means to repress stress fiber formation and EMT during TGFβ2 stimulation of lens epithelial cells. Further studies aimed at elucidating Tpm3.1 in *in vitro* human cataract model systems^34^ as well as *in vivo* cataract animal models^19^ are required to determine the therapeutic potential of targeting Tpm3.1 in cataracts. Targeting Tpms could be a promising route to prevent EMT and cataractogenesis.

## Acknowledgements

This work was funded by the National Institutes of Health (NIH) Grant R01 EY017724 to V.M.F. J.P. was supported by a fellowship from the Natural Science and Engineering Research Council of Canada. E.K. was supported by a Life Sciences Summer Institute internship from the San Diego Workforce Partnership.

## Supplemental Figures

**Figure S1.**
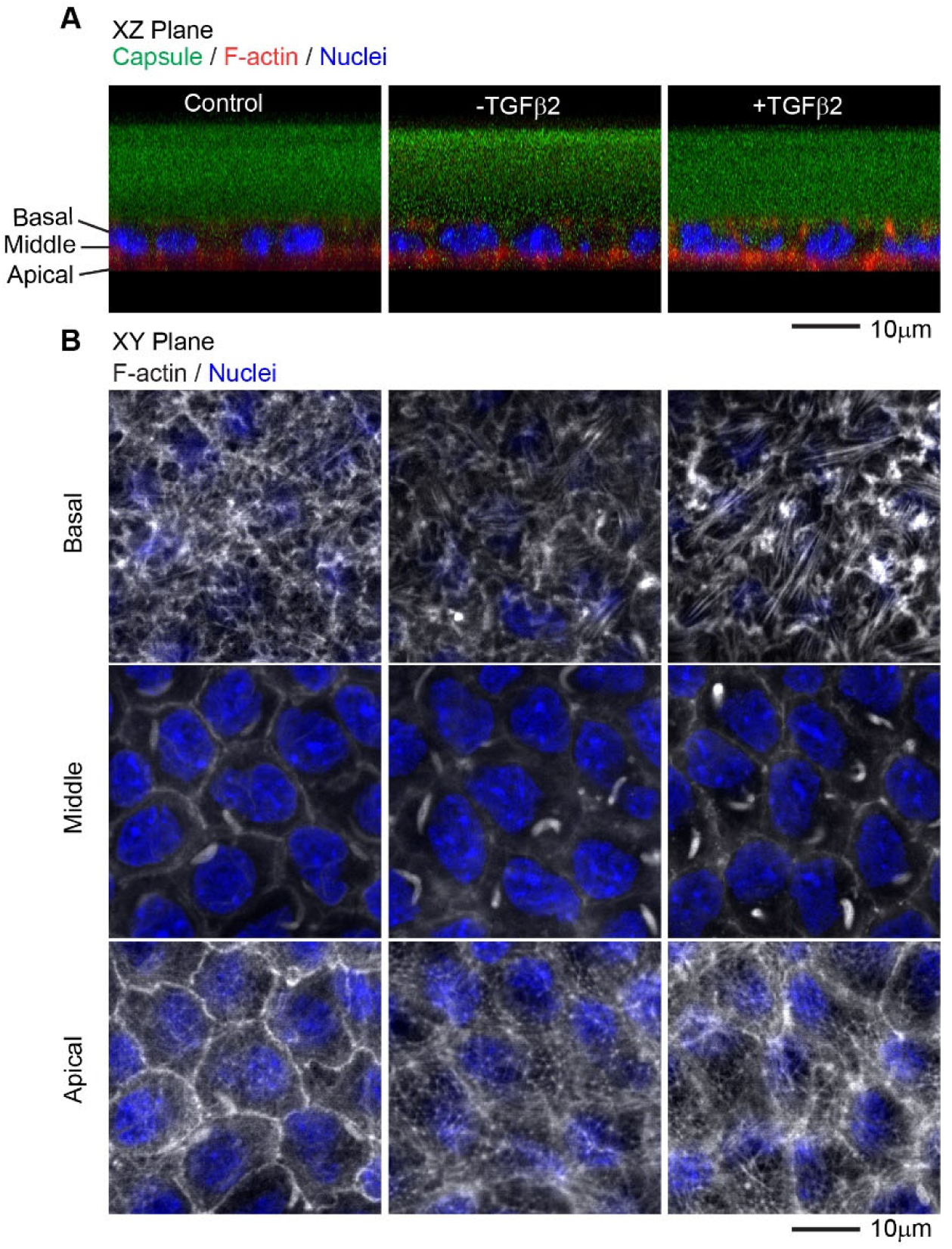
Culture of whole mouse lenses for 2 days results in stress fiber formation at the basal region of lens epithelial cells. Whole mount confocal images of lenses stained for capsule, F-actin with phalloidin, and nuclei. (A) Sagittal (*x,z*-plane view) optical sections from 3D reconstructions of confocal z-stacks showing the capsule (WGA; green) and underlying epithelial cells stained for F-actin (phalloidin; red) and nuclei (Hoecsht; blue). (B) Single (*x,y*-plane view) optical sections of basal, lateral, and apical regions of the lens showing F-actin organization (phalloidin; grayscale) and nuclei (Hoecsht; blue). At the apical and middle regions of epithelial cells in control (non-cultured) lenses, F-actin is predominantly associated with the cell-cell junctions at the membranes^35^ or in sequestered actin bundles. Additionally, F-actin is also within polygonal arrays at the apical regions. There are little to no stress fibers at the basal regions of non-cultured lenses. Culture of lenses in TGFβ2 results in slightly dimmer staining for F-actin at cell-cell junctions, but sequestered actin bundles and polygonal arrays do not appear to be affected. Most notably, TGFβ2 treatment leads to stress fiber formation at the basal regions of epithelial cells.

**Figure S2.**
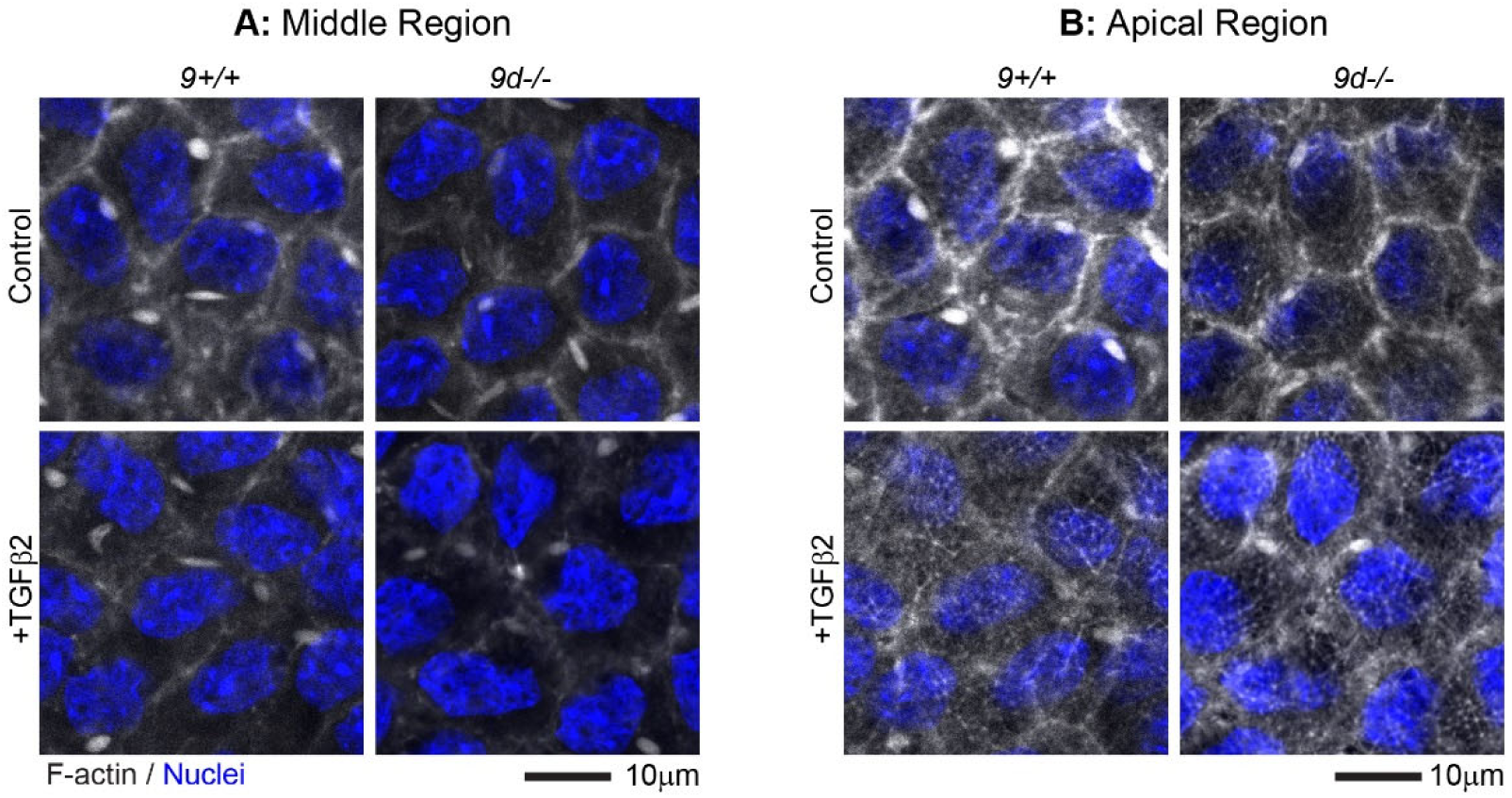
F-actin organization at middle and apical regions of the *Tpm3/Δexon9d*^*-/-*^ lenses. Whole mount images showing single (*x,y*-plane view) optical sections of (A) middle and (B) apical regions of the lens epithelium in non-cultured and lenses cultured with TGFβ2 for 2 days. In control (non-cultured) lenses, F-actin staining (phalloidin; grayscale) is similar in *Tpm3/Δexon9d*^*-/-*^ (9d-/-) lenses as compared to *Tpm3/Δexon9*^*+/+*^ (9d+/+) lenses at the middle and apical regions of lens epithelial cells. Culture of lenses in TGFβ2 results in slightly dimmer staining for F-actin at cell-cell junctions in middle and apical regions, as shown in Figure S1. This occurs in both *Tpm3/Δexon9d*^*-/-*^ (9d-/-) lenses and *Tpm3/Δexon9*^*+/+*^ (9d+/+) lenses.

